# Bats host the most virulent—but not the most dangerous—zoonotic viruses

**DOI:** 10.1101/2021.07.25.453574

**Authors:** Sarah Guth, Nardus Mollentze, Katia Renault, Daniel G. Streicker, Elisa Visher, Mike Boots, Cara E. Brook

## Abstract

Identifying virus characteristics associated with the largest public health impacts on human populations is critical to informing zoonotic risk assessments and surveillance strategies. Efforts to assess “zoonotic risk” often use trait-based analyses to identify which viral and reservoir host groups are most likely to source zoonoses but have not fully addressed how and why the impacts of zoonotic viruses vary in terms of disease severity (‘virulence’), capacity to spread within human populations (‘transmissibility’), or total human mortality (‘death burden’). We analyzed trends in human case fatality rates, transmission capacities, and total death burdens across a comprehensive dataset of mammalian and avian zoonotic viruses. Bats harbor the most virulent zoonotic viruses even when compared to birds, which alongside bats, have been hypothesized to be “special” zoonotic reservoirs due to molecular adaptations that support the physiology of flight. Reservoir host groups more closely related to humans—in particular, Primates—harbor less virulent, but more highly transmissible viruses. Importantly, disproportionately high human death burden, arguably the most important metric of zoonotic risk, is not associated with any animal reservoir, including bats. Our data demonstrate that mechanisms driving death burdens are diverse and often contradict trait-based predictions. Ultimately, total human mortality is dependent on context-specific epidemiological dynamics, which are shaped by a combination of viral traits and conditions in the animal host population and across and beyond the human-animal interface. Understanding the conditions that predict high zoonotic burden in humans will require longitudinal studies of epidemiological dynamics in wildlife and human populations.

**Significance statement:** The clear need to mitigate zoonotic risk has fueled increased viral discovery in specific reservoir host taxa. We show that a combination of viral and reservoir traits can predict zoonotic virus virulence and transmissibility in humans, supporting the hypothesis that bats harbor exceptionally virulent zoonoses. However, pandemic prevention requires thinking beyond zoonotic capacity, virulence, and transmissibility to consider collective ‘burden’ on human health. For this, viral discovery targeting specific reservoirs may be inefficient as death burden correlates with viral, not reservoir, traits, and depends on context-specific epidemiological dynamics across and beyond the human-animal interface. These findings suggest that longitudinal studies of viral dynamics in reservoir and spillover host populations may offer the most effective strategy for mitigating zoonotic risk.

## Introduction

The vast majority of human pathogens are derived from animal populations (1). In response to increasingly frequent zoonotic spillovers and their substantial public health risks (2), there has been a movement to identify the ecological systems and taxonomic groups of animals and pathogens that are most likely to source the next emerging zoonosis in the human population (3– 9). However, most of this work has centered on a binary definition of zoonotic risk—whether particular pathogens are capable of infecting humans—without considering how pathogens vary with respect to their impacts on humans after spillover. The ongoing SARS-CoV-2 pandemic has re-emphasized the reality that not all zoonoses pose risks of equal magnitude—some are exceptionally more dangerous than others due to the severity of disease they cause (‘virulence’) or their capacity to spread within human populations (‘transmissibility’), which combined, influence the total number of human deaths (‘death burden’) (10). Given the extraordinary diversity of both animal hosts and the viruses they harbor, understanding which animal and virus groups are more likely to source dangerous zoonoses is an important public health aim. Many high-profile zoonotic viruses—including Nipah and Hendra henipaviruses; Ebola filovirus; SARS, MERS, and SARS-CoV-2 coronaviruses; pandemic avian influenzas; West Nile virus; and Eastern Equine encephalitis virus—have emerged from Chiropteran (bat) or avian reservoirs (11). The high number of zoonotic viruses found in bats and birds has been attributed to their large gregarious populations, mobility, ability to colonize anthropogenic environments, and sheer species diversity (7, 11). Nonetheless, the question remains: are bat- and/or bird-borne viruses disproportionately dangerous?

A recent meta-analysis (10) found that mammalian reservoir hosts most closely related to humans harbor zoonoses of lower impact in terms of mortality relative to more phylogenetically distant hosts. These results were consistent with phylogenetic trends in virulence that have been reported in cross-species pathogen emergences in other systems (12, 13), and likely reflect mismatches in host biology, physiology, and ecology. Notably, order Chiroptera (bats)—one of the more distantly related host orders—had the highest positive effect size on case fatality rate in humans. Nevertheless, this analysis considered only directly transmitted viruses and viruses derived from mammalian hosts, despite the existence of several high-profile vector-borne and avian zoonoses (11). In particular, birds occupy a separate taxonomic class from humans—a phylogenetic distance that might correlate with heightened virulence in humans.

*In vitro* work has suggested that molecular adaptations that support the physiology of flight, a trait unique to bats among mammals, may allow bats to tolerate rapidly-replicating viruses that express heightened virulence upon emergence in less tolerant hosts such as humans (14)—thus offering a possible explanation for bat virus virulence. Bats and birds share a suite of convergent flight adaptations—both taxa are remarkably long-lived for their body size and appear to circumvent metabolic constraints on longevity through cellular pathways evolved to mitigate oxidative stress induced by flight (11). These metabolic adaptations are hypothesized to be linked to the evolution of virulent viruses in bats, but only typically discussed with respect to their effect on lifespan in birds (15). A few papers have reviewed birds’ role as special zoonotic reservoirs (11, 16), but the virulence of avian zoonoses remains largely unexplored. Nonetheless, though the most virulent zoonotic viruses may garner the most publicity, these pathogens are not necessarily the most ‘dangerous’ to human health. Rather, human health is most impacted by viruses that cause large volumes of cases and deaths (‘burden’). While some viruses such as Ebola and rabies are associated with both high case fatality rates and burden in the human population, pandemic viruses are often characterized by relatively low case fatality rates but high human transmissibility. The 2009 H1N1 influenza pandemic was estimated to have caused 60.8 million cases and more than 12,000 deaths in the United States alone with a case fatality rate of less than 1% (17), and as of July 9^th^, 2021, SARS-CoV-2 has caused over 185 million cases and 4 million deaths worldwide with a case fatality rate of just 2.2% (18). To prevent the next zoonotic pandemic, it is important to think beyond the individual measures of zoonotic capacity, virulence, and transmissibility to consider collective ‘burden’ on public health.

We apply generalized additive models (GAMs) to a comprehensive dataset of mammalian and avian zoonotic viruses to identify reservoir host and viral traits predictive of the (a) case fatality rate (CFR), (b) capacity for forward transmission, and (c) death burden induced by infections in the human population—with the goal of characterizing sources of zoonotic viruses that pose the greatest danger to global health. Our work builds on a small body of meta-analyses that have begun to explore variation in the virulence and between-human transmissibility of zoonotic viruses (4, 19–21). We provide the most thorough analysis of quantitative zoonotic virus data published to date, including the first analysis of burden and the largest sample size—with trends examined across the majority of known zoonotic viruses. We hypothesized that birds—given their capacity for flight and phylogenetic distance from humans—might rival bats for the association with the most virulent zoonotic viruses. However, we did not expect bats or birds to be responsible for the greatest burden on global health, instead anticipating high burden to be largely a function of viral traits and associated with reservoir orders that harbor less virulent, more transmissible viruses.

## Results

Drawing from existing databases (3, 7), we compiled a dataset of all mammalian and avian zoonotic virus species that met a strict definition of zoonotic—requiring a record of natural human infection confirmed by PCR or sequencing and animal-to-human directionality in transmission. Virus species linked to multiple independent reservoir groups (e.g., canine and bat rabies) or those which spillover to humans both directly from their reservoir and through bridge hosts (e.g., Nipah virus) were subdivided into separate entries for each unique transmission chain ending in spillover, creating a final dataset of 87 viruses with a total of 91 transmission chains (*SI Data and Results*, Table S1). We then applied generalized additive models (GAMs) to assess predictors (*SI Data and Results*, Table S7) of three metrics of zoonotic risk: global estimates of case fatality rates (CFRs) in humans (proxy for virulence), capacity for forward transmission within the human population ranked on a four-point scale (human transmissibility), and post-1950 cumulative death counts (death burden) (*Materials and Methods*).

### Predictors of human CFRs

In our virulence analysis, we observed a left-skewed distribution of CFRs, with 34.1% of virus species linked to no fatalities (0% CFR) and more than half (58.5%) linked to a CFR of less than 10% (Figure S1 in *SI Figures*). Bat reservoirs harbored the most virulent zoonotic viruses, contributing two thirds of the identified viruses with CFRs higher than 50%. The top selected GAM to predict global estimates of CFR in humans—across the 86 unique zoonotic transmission chains for which at least two human cases have been recorded— explained 74.7% of the deviance and included virus family, reservoir host group, bridged spillover, and vector-borne transmission (Figure 1, Table S5a in *SI Data and Results*). Consistent with previous work (10) and the hypothesis that bats are “special” zoonotic reservoirs, order Chiroptera had the largest positive effect size on CFR in humans (Figure 1b). The top selected model predicted a CFR of 65.4% for zoonotic viruses derived from order Chiroptera, representing a more than 50% increase from the next highest predicted CFR (Figure S2). Contrary to our flight hypothesis, avian reservoirs were not similarly associated with disproportionately virulent zoonoses; order Aves had a neutral effect size on human CFR that was not significant. Order Cetartiodactyla had the largest negative effect size on CFR, but notably, Cetartiodactyl hosts in our dataset included only domesticated animal species—cattle, pigs, and camels. The long coexistence of domestic animals and humans likely facilitated increased research effort for this clade, which have may have led to greater detection of low virulence zoonoses in domestic animal species. A long history of domestic animal-human coexistence may also have supported the development of preexisting human immunity to some livestock diseases, resulting in lower virulence infections.

**Figure 1.**
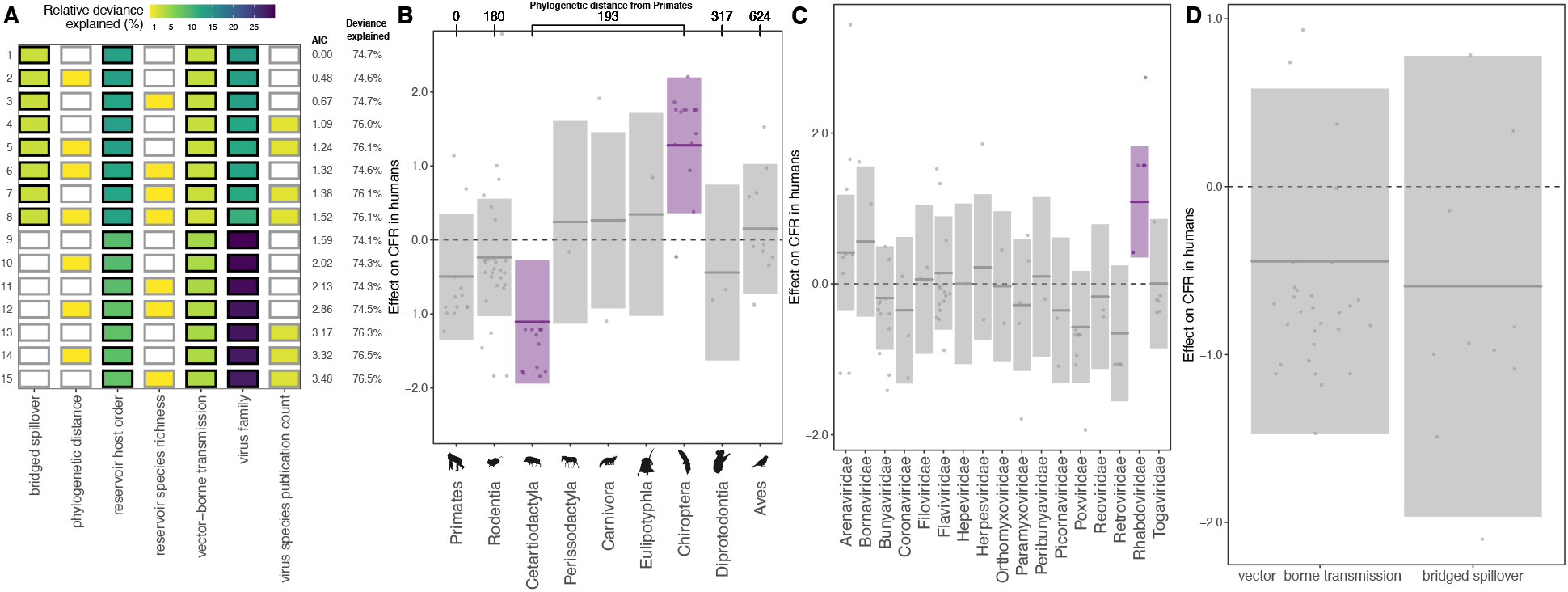
Predictors of global CFR estimates. (A) Top 15 models ranked by AIC. Rows represent individual models and columns represent predictor variables. Cells are shaded according to the proportion of deviance explained by each predictor. Cells representing predictor variables with a p-value significance level of <0.1 are outlined in black. (B-D) Effects present in the top model: reservoir host group, virus family, vector-borne transmission, and bridged spillover. Lines represent the predicted effect of the x-axis variable when all other variables are held at their median value (if numeric) or their mode (if categorical). Shaded regions indicate 95% CIs by standard error and points represent partial residuals. An effect is shaded in gray if the 95% CI crosses zero across the entire range of the predictor variable; in contrast, an effect is shaded in purple and considered “significant” if the 95% CI does not cross zero. Full model results are outlined in Table S5a in *SI Data and Results*. (B) Reservoir host groups are ordered by increasing cophenetic phylogenetic distance from Primates (in millions of years), as indicated on the top axis.

Past analyses have observed that particular viral families associate non-randomly with particular host groups (10, 22), suggesting that virus taxonomy may underlie trends in virulence across reservoir orders. For example, the high number of virulent bat-borne zoonoses (Figure S1 in *SI Figures*) may be entirely a result of the virus groups that preferentially infect bats, rather than the bats themselves. However, here, reservoir host group and virus family significantly predicted CFR within the same models (Figure 1a), indicating that both reservoir and virus taxa contributed to the observed variation in virulence. Chiroptera had the highest positive effect size on CFR despite being associated with virus families that ranged from the most (Rhabdoviridae) to least (Coronaviridae) virulent (Figure 1c). Removing the 100% fatal lyssaviruses (n=5) from the dataset resulted in large reductions in the CFR predicted for bat-borne zoonoses (Figure S4), though order Chiroptera still had the highest and most significant positive effect size on CFR (Figure S2 in *SI Figures*, Table S6a in *SI Data and Results*).

Previous work has demonstrated a positive correlation between reservoir host phylogenetic distance from humans and the case fatality rates of zoonoses derived from those reservoirs (10); in our analysis, however, reservoir host group phylogenetic distance from Primates was not correlated with CFR, dropping entirely from the top ranked model and not ranking significantly in any of the top 15 selected models (Figure 1a). The combined effect of reservoir host group and virus family as predictor variables in the same model likely overwhelmed any correlation between host phylogeny and CFR, particularly given the lack of granularity in our phylogenetic distance variable, based on a time-scaled phylogeny, which produced only six unique distance values across nine host groups, with Chiroptera and four of the other mammalian orders clustering at a single distance level (*Materials and Methods*). Nevertheless, trends in effect size on CFR (Figure 1b) and predicted CFR (Figure S2) across reservoir host groups suggest that, in general, virulence increases with phylogenetic distance, but this positive correlation may collapse at “extreme” distances.

To test whether these results held across a larger sample size, we ran a CFR analysis that included viruses that met a more lenient definition of zoonotic—specifically, viruses with only serological evidence of infection in humans, viruses that have only caused human infections in laboratory settings, and viruses for which only one human case has been recorded—increasing our dataset to 119 virus species with a total of 123 unique zoonotic transmission chains (Figure S5 in *SI Figures*, Table S6b in *SI Data and Results*). This supplementary analysis echoed the results from our first analysis of global CFR estimates—both reservoir and virus taxonomy contributed to the observed variation in CFR (Figure S5a in *SI Figures*); and Chiroptera had the highest positive effect size on CFR, whereas Aves had a neutral nonsignificant effect (Figure S5b in *SI Figures*).

To assess whether CFR trends might be influenced by health care differences among the virus’ differing geographic ranges, we tested whether Gross Domestic Product per capita (GDPPC) significantly predicted country-specific CFR estimates—calculated from death and case counts in countries that have reported the largest outbreaks of each given virus species, with up to three country estimates for each species for a total of 119 estimates across the 86 unique zoonotic transmission chains. First, we modeled all 119 country-specific CFR estimates separately to test whether GDPPC predicts country-level variation in CFR (Figure S6 in *SI Figures*, Table S6c in *SI Data and Results*). Although significant, GDPPC explained a low percentage of the deviance (Figure S6a in *SI Figures*), and wide confidence intervals indicated uncertainty in trends (Figure S6d in *SI Figures*). To gage whether variation in GDPPC among virus’ geographic ranges might bias the trends in global CFR estimates observed in the Figure 1 models, we then modeled GDPPC and country CFR estimates aggregated at the level of the 86 unique zoonotic transmission chains (Figure S7 in *SI Figures*, Table S7d in *SI Data and Results*).

GDPPC was not significant in any of the top models, often dropping entirely during model selection (Figure S7a in *SI Figures*), suggesting that health care differences among the virus’ geographic ranges most likely do not bias Figure 1 trends. Nevertheless, as with the supplementary analysis presented in Figure S5, both analyses of the country CFR estimates echoed all key results presented in Figure 1.

### Predictors of transmissibility within human populations

We found that most zoonotic viruses (72.1%) have not been reported to transmit within the human population following spillover (i.e., transmissibility rank = 1, or R0 = 0) (Figure S8). Only 15.1% of virus species had demonstrated capacity for endemic transmission among humans, of which the majority (61.5%) were sourced from Primates. The top selected GAM to predict the ordinal rank of transmissibility within human populations—across the 86 unique zoonotic transmission chains for which at least two human cases have been recorded—explained 56.7% of the deviance and included virus family, the phylogenetic distance between each virus’ reservoir host group and Primates, vector-borne transmission, and the virus species publication count (Figure 2, Table S5b in *SI Data and Results*). Transmissibility declined with phylogenetic distance from Primates, but the estimated trend was highly uncertain (Figure 2c). We therefore reran the analysis with reservoir host group as the only host taxonomic predictor (excluding the phylogenetic distance variable). This analysis identified Primates as the only host order significantly associated with heightened transmissibility in humans, suggesting that this group is the primary driver of the phylogenetic trend observed in the top selected model (Figure S9a in *SI Figures*, Table S6c in *SI Data and Results*).

**Figure 2.**
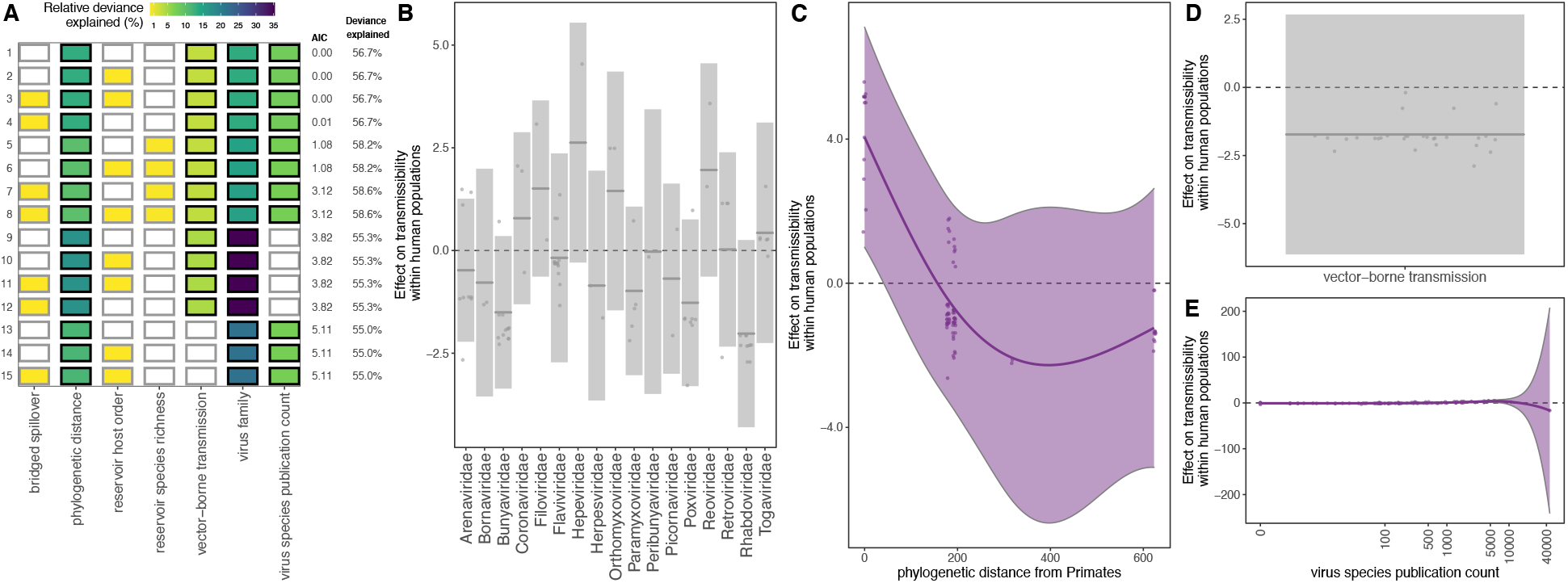
Predictors of capacity for forward transmission within the human population following zoonotic spillover. (A) Top 15 models ranked by AIC. Rows represent individual models and columns represent predictor variables. Cells are shaded according to the proportion of deviance explained by each predictor. Cells representing predictor variables with a p-value significance level of <0.1 are outlined in black and otherwise outlined in gray. (B-E) Effects present in the top model: virus family, reservoir group phylogenetic distance from Primates, vector-borne transmission, and virus species publication count. Lines represent the predicted effect of the x-axis variable when all other variables are held at their median value (if numeric) or their mode (if categorical). Shaded regions indicate 95% CIs by standard error and points represent partial residuals. An effect is shaded in gray if the 95% CI crosses zero across the entire range of the predictor variable; in contrast, an effect is shaded in purple and considered “significant” if the 95% CI does not cross zero. Full model results are outlined in Table S5b in *SI Data and Results*.

Evolution of virulence theory typically assumes a tradeoff between virulence (death rate due to infection) and transmission rate on the basis that while high within-host growth rates increase infectiousness, they also increase damage to the host, increasing virulence and thus shortening the infectious period and reducing opportunities for future transmission (23, 24). Critically, CFR is not equivalent to virulence, but instead, a proxy that can be reliably quantified. As defined by Day 2002 (25), CFR is a function of both pathogen virulence (*α*) and clearance rate (*σ*), in which *CFR* = *α*/(*α* + *σ*). Thus, virulent pathogens (high *α*) with high clearance rates (high *σ*)—e.g., acute, short-lived infections such as Chikungunya virus (26)—could produce low CFRs. In contrast, less virulent pathogens (low *α*) with low clearance rates (low *σ*)—e.g., persistent infections such as HIV (27)—could produce high CFRs. Nevertheless, in our data, we observed a relationship between CFR and transmissibility in humans that roughly supports the fundamental theoretical tradeoff between virulence and transmission rate (Figure S10 in *SI Figures*). Viruses causing the highest CFRs in humans (>75% CFR) clustered in the lower right corner with the lowest capacity for forward transmission in the human population, implying maladaptive virulence. Conversely, the least virulent viruses (0% CFR) clustered at either the lowest transmission capacity—likely indicative of poor compatibility with humans—or the highest transmission capacity—suggesting transmission uninhibited by virulence.

### Predictors of post-1950 death burden in the human population

For our death burden analysis, we modeled the total number of deaths resulting from a given zoonosis recorded worldwide since 1950 (and up until March 7^th^, 2021). In cases where our death count could only begin after 1950, either because a zoonosis first emerged in humans after 1950 or because reliable death records were only available for a subset of the timeline, we standardized analyses by including an offset for the number of years over which the death counts were recorded. The raw death count distribution was highly left-skewed, with 39.5% of virus species linked to 0 deaths and more than half (62.7%) linked to fewer than 50 deaths (Figure S11 in *SI Figures*). We observed significant overdispersion in death counts, even when standardized by the number of years over which the deaths were recorded, with deaths per year ranging from zero to almost 2 million for SARS-CoV-2. Just two viral predictors—virus family and species publication count—explained most of the variation in death burden among the 91 zoonotic transmission chains across all the top GAMs (Figure S12a in *SI Figures*). Host predictors explained a very low percentage of the variation in death burden across all the top selected models, often dropping entirely during term selection. Virus species publication count tempered virus family effects (Figure S12c in *SI Figures*) because virus species with high death burdens were also associated with high publication counts, likely because high death burdens motivate increased research efforts. In contrast, there was little evidence that poorly studied viruses had unusually low death burdens, implying that a lack of diagnostic effort is not a major driver of low death burdens in our data (Figure S12c in *SI Figures*). After excluding the virus species publication predictor, we found that Coronaviridae, Orthomyxoviridae and Rhabdoviridae had the highest positive effect sizes on death burden, driven by, respectively, the SARS-CoVs, the Influenza A transmission chains, and Rabies virus (Figure 3b, Table S5c in *SI Data and Results*). With virus publication count removed, the top four models included two reservoir traits—phylogenetic distance from Primates and species richness—as significant predictors. Reservoir groups most closely related to Primates were associated with heightened death burdens relative to more distantly related reservoirs, consistent with results from our transmissibility analyses that indicated that reservoirs most closely related to Primates harbored more transmissible viruses (Figure 3c). Reservoir species richness positively correlated with death burden, as we would expect given that species richness has been found to correlate with the number of viruses associated with a given reservoir order (Figure 3d) (7). However, both reservoir predictors explained a small fraction of the variation in death burden relative to virus family, confirming that death burden is largely a function of viral traits (Figure 3a).

**Figure 3.**
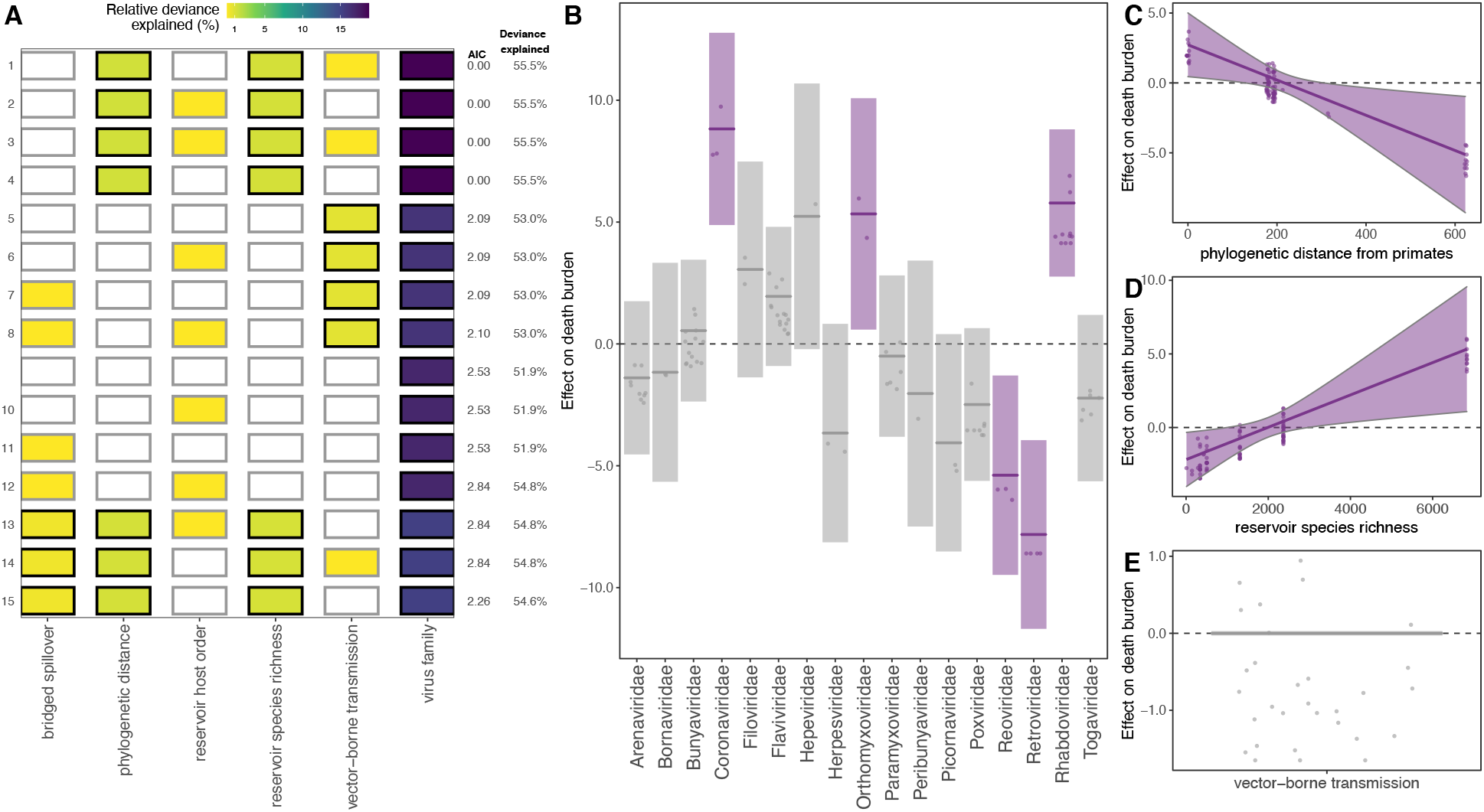
Predictors of post-1950 death burden, excluding the virus species publication count predictor. See Figure S12 in *SI Figures* for inclusion. (A) Top 15 models ranked by AIC. Rows represent individual models and columns represent predictor variables. Cells are shaded according to the proportion of deviance explained by each predictor. Cells representing predictor variables with a p-value significance level of <0.1 are outlined in black and otherwise outlined in gray. (B-D) Effects present in the top model: virus family, reservoir group phylogenetic distance from Primates, reservoir group species richness, and vector-borne transmission. Lines represent the predicted effect of the x-axis variable when all other variables are held at their median value (if numeric) or their mode (if categorical). Shaded regions indicate 95% Cis by standard error and points represent partial residuals. An effect is shaded in gray if the 95% CI crosses zero across the entire range of the predictor variable; in contrast, an effect is shaded in purple and considered “significant” if the 95% CI does not cross zero. Full model results are outlined in Table S5c in *SI Data and Results*.

While some reservoir groups—bats, primates, rodents, and birds—have sourced more high burden viruses than others (Figure 4a), both our model results and raw data suggested that high burden viruses appeared to be function of viral traits, not the reservoirs themselves. No single reservoir stood out as a consistent source of high burden viruses, with every reservoir that harbors high burden viruses also harboring substantially more viruses that cluster at the lowest death burdens (Figure 4a). This was not the case for virus family (Figure 4b) or primary transmission route (Figure 4c); Coronaviridae and Orthomyxoviridae and a respiratory transmission route were associated only with high burden zoonotic viruses. In general, the viruses linked to the lowest death burdens were associated with the lowest transmission capacity. As a deviation from this trend, Primates—which our models indicate harbor the most transmissible, but generally less virulent zoonotic viruses—harbored several highly transmissible viruses with low death burdens (Figure 4a).

**Figure 4.**
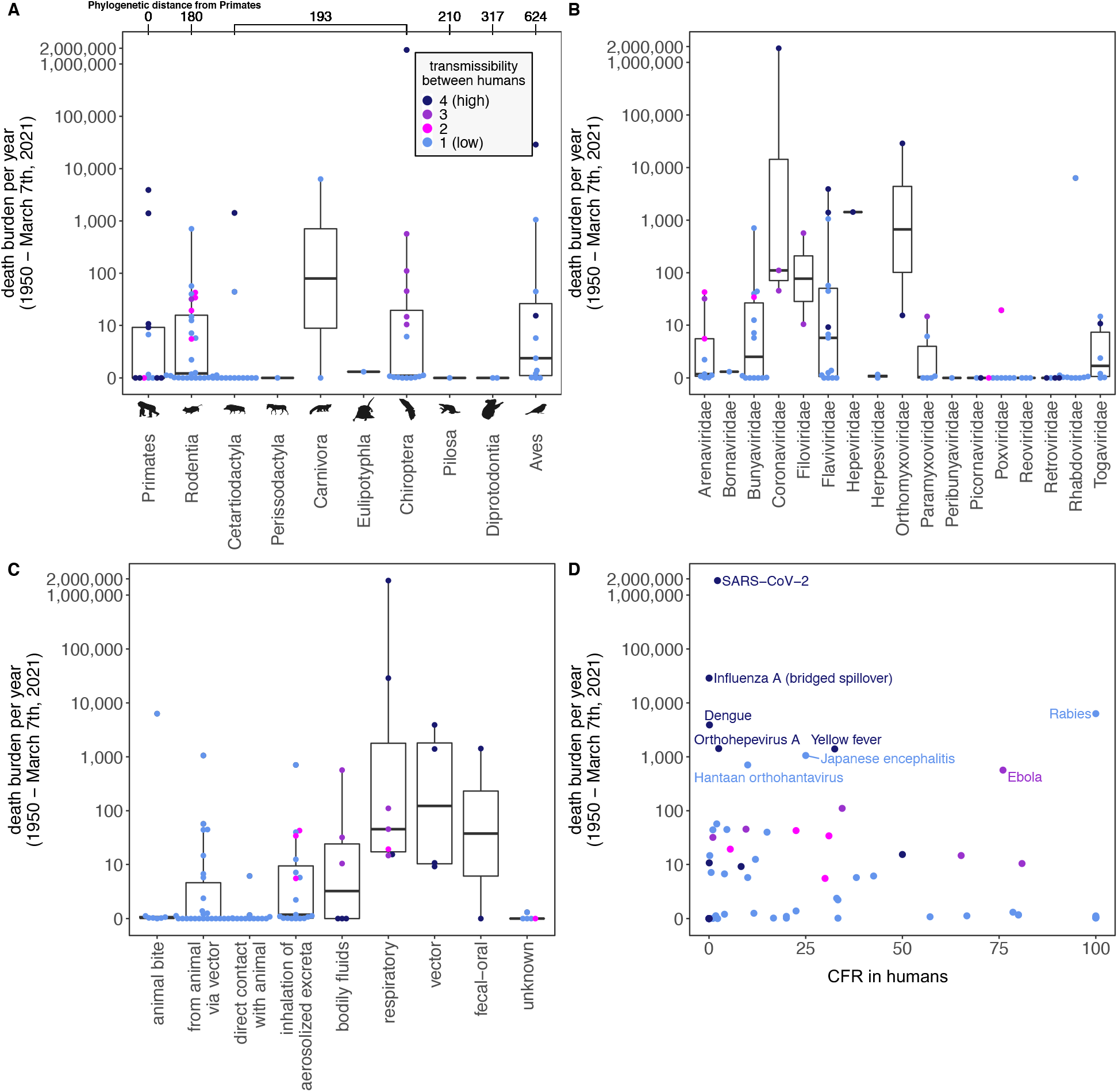
Death burden per year (cumulative post-1950 death counts divided by the length of reporting time), grouped by (A) reservoir host group, (B) virus family, (C) primary transmission route, and (D) CFR in humans. Colors indicate transmissibility between humans, with “1” indicating the lowest level of transmission (i.e., no recorded forward transmission in human population post-spillover) and “4” indicating the highest level of transmission (i.e., record of endemic transmission in human populations post-spillover). (A) Reservoir host groups are ordered by increasing cophenetic phylogenetic distance from Primates (in millions of years), as indicated on the top axis.

The highest death burdens were overall associated with zoonotic viruses that are less virulent but highly transmissible in human populations (Figure 4d). Respiratory pathogens with capacity for human-to-human transmission have often incurred massive burdens over short timeframes as a result of rare, but catastrophic spillover events that spark widespread transmission in humans. Critically, while our dataset included only six viruses with respiratory droplets as a primary transmission route—SARS-CoV-1, SARS-CoV-2, MERS CoV, Influenza A, Nipah, and Monkeypox—these viruses accounted for more than 85.9% of the deaths recorded for the 86 viruses in our death burden analysis, highlighting respiratory transmission as a high-risk zoonotic trait. However, these data were derived from a notably small sample size, as three of the six respiratory viruses have caused only a single major epidemic. There was also substantial variation among these respiratory viruses, with the death burdens associated with SARS-CoV-1 and SARS-CoV-2 differing by more than 2.5 million.

Additionally, several outliers demonstrated that capacity for forward transmission in human populations does not always predict death burden; it is critical to also consider epidemiological dynamics across and beyond the human-animal interface. Less transmissible viruses can accumulate large death burdens over many small, but frequent spillovers, particularly in systems in which humans regularly interact with animal reservoirs. Rabies, Hantaan (HTNV), and Japanese Encephalitis viruses have been associated with some of the highest death burdens induced by viral zoonosis despite lacking forward transmission in human populations (Figure 4d). This is likely because these viruses spill over to humans from animal host populations that live amongst human communities—Rabies burden is largely driven by spillover from endemic circulation in domestic dogs (28), HTNV spills over from striped field mouse (*Apodemus agrarius*) populations that inhabit agricultural fields (29), and Japanese encephalitis is amplified via domesticated pigs (30). Outbreaks in these spillover host populations source human infections that are dead ends for further transmission but add up to large numbers. Emphasizing the importance of understanding system-specific dynamics, HTNV had a death burden more than 18 times greater than the combined death burden of all ten other rodent-borne hantaviruses in our dataset, most likely because other rodent reservoirs of hantaviruses tend to overlap less with human populations (29). Furthermore, zoonotic viruses that have historically been low burden pathogens can “unexpectedly” cause high death burdens in the case of virus evolution or unique epidemiological circumstances (31). For example, Ebola virus first emerged in humans in 1976, causing deadly, but local outbreaks up until late 2013, when suddenly, emergence in a region with dense and interconnected human populations, coupled with virus adaptation (32), allowed an Ebola spillover event to spark a transnational epidemic that in just 2 years, caused more than 6.5 times the total number of deaths recorded from 1976-2013 (31, 33). These outliers suggest that understanding epidemiological dynamics—within wildlife populations and across and beyond the human-animal interface—in specific systems is a critical component of predicting death burden and consequently, danger to human health.

## Discussion

A key insight from our work is that bats harbor the most virulent zoonotic viruses relative to other mammalian and avian reservoirs (Figure S1 in *SI Figures*). Given that birds represent the only other flying vertebrates and that flight adaptations are hypothesized to influence viral virulence in bats (11), we expected avian viruses to similarly be associated with heightened CFRs in humans. However, we found that only order Chiroptera had an exceptionally high positive effect size on CFR in humans, while Aves had a neutral nonsignificant effect. It is of course possible that we observed this association between Chiroptera and high CFRs in part because low virulence zoonotic viruses have gone undetected in bat reservoirs; however, other poorly studied reservoirs are not comparably associated with heightened virulence, suggesting that detection bias cannot explain our results. Like CFR, transmissibility in humans was also correlated with reservoir traits, but in this case, Primates—the reservoir group most closely related to humans—sourced the zoonotic viruses with the highest capacities for forward transmission in human populations. While a combination of both virus and reservoir taxonomy predicted virulence and transmissibility, death burden did not correlate with any reservoir group and instead, was a function of viral traits. Nevertheless, our data indicated that mechanisms driving high death burdens are diverse and often contradict trait-based predictions. Several high-profile zoonotic viruses linked to significantly higher death burdens than we would expect based on their capacity for forward transmission in the human population (Figure 4d), suggesting that death burden is highly dependent on both the contact rate at the human-animal interface and epidemiological dynamics within the human population—factors which are not fully captured by the broad explanatory variables considered in trait-based analyses.

The surprisingly low virulence of avian zoonotic viruses in contrast to bat-borne viruses may reflect the extreme phylogenetic distance that separates birds from Primates. In our previous analysis, we found that mammalian reservoir hosts most closely related to humans harbor less virulent zoonotic viruses relative to more distantly related mammalian hosts such as bats (10). This positive correlation between reservoir phylogenetic distance from humans and viral virulence is consistent with trends that have been reported in cross-species pathogen emergences in other systems (10, 12, 13), and likely reflects maladaptive virulence resulting from mismatches in host biology, physiology, and ecology. Clearly, while bats are distantly related to humans, they are still mammals, whereas birds occupy a separate taxonomic class. It is likely that the positive correlation between phylogenetic distance and virulence collapses at distances beyond mammals, because viruses are expected to have a limited capacity to replicate in host environments that are very different from that of their reservoir, leading to ‘non-host resistance’ (34, 35). Phylogenetic distance dropped from all CFR models likely due to a lack of granularity in our phylogenetic distance data, which described reservoir host cophenetic distance from Primates on a time-scaled phylogeny (7), producing only six unique distance values across all of the reservoir groups in our database. Trends across reservoir host groups overall support the hypothesis that the positive correlation between phylogenetic distance and virulence collapses at “extreme” distances. Nevertheless, more studies are needed to parse the effect of phylogenetic distance on virulence trends in animal-to-human spillovers. The time-scaled phylogeny represents the only available phylogeny that includes both mammals and birds. Future studies would benefit from developing additional phylogenies of mammalian and avian reservoirs, which prioritize immunological or physiological traits that may more accurately proxy virologically relevant differences in host environments.

Chiroptera represented an outlier among distantly related reservoirs, with an undeniably positive effect size on CFR more than triple that recovered for any other mammalian order. Consistent with the hypothesis that bats represent a ‘special’ viral reservoir (36), the order Chiroptera does appear to harbor zoonotic viruses that are uniquely virulent upon spillover to humans, even when considering virulence effects that might be attributed to their phylogenetic distance from Primates. In bats, flight adaptations have been linked to viral tolerance, which previous work suggests may select for high growth rate viruses that could manifest as virulent upon emergence in less tolerant hosts such as humans (14). Notably, bats experience limited morbidity or mortality from intracellular infections with only a few known exceptions (36–39). Conversely, while birds harbor several zoonotic viruses that are virulent in humans such as Highly Pathogenic Avian Influenza (HPAI), West Nile, and Equine Encephalitis viruses, only some avian species are tolerant of these infections—many avian species experience morbidity and mortality (40). Bats and birds are expected to experience similar selective pressures from flight—they have been found to incur comparable energetic costs while flying, despite different forms and physiologies (15, 41). However, the two taxonomic groups, within disparate vertebrate classes, may have responded differently to these selective pressures. Specifically, there is a possibility that bats evolved cellular pathways that protect against both aging and immunopathology, whereas birds evolved pathways that only protect against aging. For example, bats have been found to host a suite of cellular-level anti-inflammatory adaptations—including enhanced cellular autophagy and downregulated signaling pathways linked to the induction of inflammatory antiviral defenses—which may both mitigate cellular damage induced by bat metabolism and inhibit immunopathology incurred upon viral infection (36, 42–46). On the other hand, birds may rely primarily on systemic antioxidant responses (47), which mitigate oxidative stress, but do not interact so tightly with cellular-level processes that impact viral pathology. Critically, birds appear to be missing anti-inflammatory protein tristetraprolin (TTP) (48), and immunopathology is often the cause of death in birds that die from viral infections such as HPAI and West Nile virus (40). Differences between mammalian and avian immune systems may additionally play a role in their differing infection outcomes. The immune system is broadly conserved in amniotes, but some avian immunological features diverge from those of bats and other mammals: notably, birds lack lymph nodes and instead develop B cells in a specialized lymphoid organ, the bursa of Fabricius; have heterophil in their white blood cells as opposed to neutrophil; and produce only three classes of immunoglobulin in contrast to the five produced by mammals (11). Nevertheless, the differing effects of Chiropteran and avian metabolic adaptations on viral tolerance and viral evolution remain largely uncharacterized and more basic research in this field is needed (49).

We found that both reservoir host and virus taxonomy predict the virulence and transmissibility of a virus in the secondary human host, consistent with the expectation that a virus evolves virulence to maximize reproduction in its reservoir population (50). The optimal balance between virulence and transmission depends on how the reservoir host population responds to the virus (the ‘host selective pressure’), which is determined by the ecological, physiological, and biological traits of the reservoir. While we identified “special” reservoirs of virulent and transmissible zoonotic viruses, we found that the human death burden incurred by viral zoonoses does not correlate with any one reservoir host order, including bats, and instead, is a function of viral traits. Our data demonstrate that mechanisms driving high death burdens are diverse and often contradict trait-based predictions. High death burdens have resulted from rare spillover events of highly transmissible viruses that spread widely in the human population; small, but frequent spillovers of the least transmissible viruses; and historically low-burden pathogens that take off given the right ecological and evolutionary conditions. This suggests that ultimately, death burden depends on epidemiological circumstances, which should be shaped, not by reservoir host traits, but by a combination of viral traits and conditions in the animal host population and across and beyond the human-animal interface. Notably, the pandemic spread of SARS-CoV-2 can be attributed to its highly effective respiratory transmission between humans, a trait linked to its identity within Coronaviridae, rather than its bat origins (indeed, CoVs demonstrate gastrointestinal tropism in bat reservoirs) (51).

Over the course of the last decade, a significant amount of funding and research effort has been dedicated to identifying correlates of zoonotic risk, often with a long-term aspiration of identifying ways to anticipate and prevent emerging zoonoses in the future (52–54). This research increasingly prioritizes viral discovery over longitudinal studies of epidemiological dynamics and targets animal populations such as bats that have been identified as key zoonotic reservoirs. While our analysis corroborates the hypothesis that bats are a ‘special’ reservoir for virulent zoonotic viruses, we also demonstrate that viral traits—not bat reservoirs—pose the greatest danger to human health. We argue that burden, which does not correlate with any animal reservoir and instead appears to be a function of transmission conditions to and within the human population, more correctly approximates “danger” to human health than does virus virulence. While reservoir and viral traits can predict zoonotic capacity, virulence, and transmissibility, death burden is dependent on system-specific epidemiological dynamics, which are shaped by a combination of viral traits and conditions in the animal host population and across and beyond the human-animal interface. Thus, understanding and controlling the mechanisms that drive high death burdens in humans—high rates of human-animal contact and/or epidemiological dynamics in the human population that allow discrete spillover events to trigger human epidemics— requires longitudinal surveillance of specific zoonotic or potentially zoonotic viruses in both animal and human populations. There is a pressing need for more longitudinal studies of transmission dynamics in human and wildlife populations to better understand and prevent the epidemiological conditions that cultivate the most dangerous cases of zoonotic viral emergence.

## Materials and Methods

### Constructing the database

We curated a comprehensive database of mammalian and avian zoonotic viruses—and the taxonomic orders of the reservoir hosts from which they were derived—published by Mollentze et al. 2020 (7). Using the information provided in that database and supplementing with literature searches, we extracted viruses that met a strict definition of zoonotic, requiring at least one published human infection in which the virus species was confirmed by PCR, sequencing, or isolation as well as evidence of animal-to-human directionality in transmission. We excluded six viruses (Table S2 in *SI Data and Results*) that have only caused human infections in laboratory settings. We additionally did not include viruses such as HIV (55) and HCoV-299E (56) that have zoonotic origins, but have maintained separate, genetically distinct human transmission cycles since before 1950 (Table S3 in *SI Data and Results*). We excluded such viruses for several reasons: precise death and case count records are sparse pre-1950; viruses that have circulated within the human population for centuries or decades often have unconfirmed or disputed origins; and over long timescales, viral evolution in the human population is expected to muddle any relationship between zoonotic history and dynamics in the human population (31). With this strict inclusion criteria, we compiled 87 unique virus species (Table S1 in *SI Data and Results*). Each virus species was associated with one reservoir host order, with the exception of Rabies virus and Mammalian 1 orthobornavirus, which are both known to be maintained by two distinct nonhuman animal reservoir orders in independent transmission cycles (7).

For each virus-reservoir association, we collected both human case fatality rate (CFR) as a proxy for virulence, and the cumulative global death count as a proxy for burden on the human population. For CFR, we collected two estimates. First, we recorded existing estimates of global CFRs from the literature, calculating averages when ranges were reported. Second, for each virus species, we calculated up to three country-specific CFRs from death and case counts in countries that have reported the largest outbreaks of that virus—when available, using data that spanned multiple outbreaks and/or years to maximize sample size and accuracy. We expected that global CFR estimates would be more precise approximations of virulence, while country-specific CFR reports would allow us to assess and account for potentially confounding effects of regional differences in health care and overall infrastructure. For our death burden response variable, we collected the total number of deaths recorded across the world since 1950. In many cases, our death count began after 1950, either because a zoonosis first emerged in humans after 1950 or reliable death records were only available for a subset of the timeline. To standardize, we added a variable for the number of years over which death counts were recorded to use as an offset in our models. Death and case counts were derived, when available, from the Global Infectious Diseases and Epidemiology Network (GIDEON) (57)—which contains outbreak data from case reports, government agencies, and published literature records—and supplemented with literature searches. All variable descriptions are provided in Table S4 in *SI Data and Results*.

We additionally ranked each zoonosis’ capacity for transmission within human populations—a correlate of R0—on a four-point scale (10). We assigned a human transmissibility level of “1” to viruses for which forward transmission in human populations post-spillover had not been recorded; “2” to viruses for which forward transmission in humans had been recorded but was described as atypical; “3” to viruses for which transmission within human populations had occurred regularly but was restricted to self-limiting outbreaks; and “4” to viruses for which endemic human transmission had been reported.

Recording death and case data from laboratory-confirmed outbreaks in the literature required maintaining a strict definition of zoonotic, excluding some viruses that have been included in previous meta-analyses (3, 7, 19). We compiled excluded viruses that met looser inclusion criteria—specifically, seven viruses that have only caused human infections in laboratory settings and 25 viruses that lacked molecular confirmation of infection of humans, but still had serological evidence of infection in humans—in a supplementary database (Table S2 in *SI Data and Results*). Viruses included in previous meta-analyses that met neither our loose nor strict inclusion criteria are outlined in Table S3 in *SI Data and Results*.

Drawing from previously published databases (3, 7, 10), we collected seven variables (*SI Data and Results*, Table S7) that we hypothesized might predict observed variation in human CFR, capacity for transmission within human populations, and death burden. Given published correlations between phylogenetic distance and virulence in cross-species spillovers (10, 12, 13, 58, 59), we included the reservoir host group cophenetic distance from Primates. We calculated this distance variable using a composite time-scaled phylogeny of the mean divergence dates for all reservoir clades, as presented in the TimeTree database (7, 60). In our prior analysis (10), phylogenetic distance values were derived from a phylogenetic tree of mammalian cytochrome *b* sequences (3, 61, 62), which captured significantly more variation between host orders. The time-scaled phylogeny used in this analysis produced only six unique distance values across all reservoir groups in our database but represented the only available phylogeny that included both mammals and birds. We considered both reservoir host and virus taxonomy, recording host order and virus family. However, only ten avian zoonoses were distributed across several avian reservoir host orders. To test our hypotheses regarding avian zoonoses, we addressed this small sample size by aggregating avian reservoir orders into a single “Aves” group, while maintaining separate host orders for the mammalian reservoirs. Given that the number of zoonoses harbored by a reservoir group appears to correlate with species diversity within that group (7), we hypothesized that species diversity might influence reservoir effect size on CFR in humans; thus, we included reservoir species richness, which we derived from the Catalogue of Life using version 0.9.6 of the taxize library in R (7, 63), taking the sum of values across bird orders for the Aves reservoir group. If increasing a reservoir group’s total number of zoonotic viruses also increases their number of virulent zoonoses, reservoir species richness might inflate the mean CFR of zoonotic viruses harbored by species rich reservoir groups—or alternatively, given that most zoonotic viruses have low CFRs in humans, species richness might instead reduce the mean CFR associated with these reservoirs. Nevertheless, we expected that higher numbers of zoonotic virus species would inflate the total death burdens associated with species rich reservoir groups. We defined a “spillover type” variable to account for the zoonotic transmission chain of each virus, distinguishing between zoonoses that jump into humans directly from the reservoir population and those that spillover to humans from bridge hosts (10). While the majority of zoonoses were linked to single zoonotic transmission chains, there were a few exceptions with both “direct” and “bridged” spillover. For example, zoonotic Influenza A virus and Nipah virus (64, 65) have spilled over into the human population directly from their avian and bat reservoirs, respectively, as well as from domestic pig bridge host populations. In such cases, each spillover type (i.e., transmission chain) was entered separately in the database. We included an additional binary variable that identified whether viruses were vector-borne, as both theory (23) and previous meta-analyses (19, 20) have suggested a relationship between vector-borne transmission and virulence. Finally, as has been done in other similar meta-analyses, we included virus species publication count to account for any potential publication bias (3, 10, 59).

To pair with our country-specific CFR data, we collected an eighth predictor variable— gross domestic product per capita (GDPPC)—as a proxy for geographical differences in the quality of health care and epidemiological control measures.

We additionally collected, for each virus species, the transmission route that contributes the majority of human infections, extending data published by Brierley et al. (19). We then assessed trends in death burden across transmission types, hypothesizing that density-dependent transmission, as characteristic of transmission via respiratory droplets, would be associated with the highest death burdens in human populations.

### Statistical analysis

Given the non-normal distribution of our data, expected nonlinear relationships, and nested data structures within our predictor variables (66), we applied generalized additive models (GAMs) in the mgcv package in R (67) to assess predictors of CFR, transmissibility, and death burden in human populations. Rather than manually specifying higher order polynomial functions, GAMs permit the use of smooth functions to capture nonlinear relationships between response and predictor variables (66, 67). We fit continuous variables (i.e., reservoir group species richness and phylogenetic distance from Primates, and virus species publication count) as smoothed effects, and all binary (i.e., vector-borne status and spillover type) and categorical (i.e., reservoir order and virus family) variables as random effects. For variable selection, we ran all possible model combinations, ranked by AIC, and selected the models with the lowest AIC values.

We first asked, *which reservoir host and virus types are associated with elevated CFRs in human populations following spillover?* We constructed GAMs in the beta regression family to query the predictive capacity of our predictor variables (*SI Data and Results*, Table S7) on CFR in humans. We compressed our CFR range to the beta distribution interval (0,1) by applying the recommended data transformation *y*” = [*y*′(*N* − 1) + 1/2]*N*, where *N* is the sample size (68, 69). We modeled all 119 country-specific CFR estimates separately to test whether GDPPC predicts country-level variation in CFR (Table S6c in *SI Data and Results*). To gage whether variation in GDPPC among virus’ geographic ranges might confound the trends in global CFR estimates, we then modeled GDPPC and CFR estimates aggregated at the level of the 86 unique zoonotic transmission chains (Table S6d in *SI Data and Results*). For this second model, we calculated a composite GDPPC for each aggregated CFR statistic by weighting each country’s GDPPC by the proportion of cases in the CFR calculation that were recorded in each country and summing the weighted GDPPCs. We then modeled the global CFR estimates, which were not tied to any specific system. For all CFR analyses, we modeled unique zoonotic transmission chains—which we defined as unique reservoir orders and spillover type combinations per virus. As a result, zoonoses with a single reservoir host order and spillover type were modeled as a single CFR entry, while those with multiple reservoir orders and/or spillover types (e.g., Influenza A and Nipah viruses) were modeled as multiple CFR entries. We excluded five viruses for which only one human case has been recorded (Table S1 in *SI Data and Results*), deciding that we could not accurately represent a single observation as a CFR. Our final GAM analysis included 82 unique virus species with a total of 86 unique zoonotic transmission chains (Table S5a in *SI Data and Results*).

Our strict definition of zoonotic status and inclusion criteria reduced our sample size. To assess whether our observed trends held across a larger sample of zoonotic viruses, we ran an additional GAM analysis of global CFR estimates that included viruses with only serological evidence of infection in humans, viruses that have only caused human infections in laboratory settings, and viruses for which only one human case has been recorded. This supplementary GAM analysis included 119 unique virus species with a total of 123 unique zoonotic transmission chains (Table S6b in *SI Data and Results*).

We next asked, *which reservoir host and virus types are associated with elevated capacity for transmission within human populations?* We constructed a GAM in the ‘ocat’ (‘ordered categorical data’) family to query the predictive capacity of our predictor variables on transmissibility, defining the vector of categorical cut points, *θ*, to match our four-point ranking scale (*θ =* 1,2,3,4). We again excluded the five viruses for which only one human case has been recorded (Table S1 in *SI Data and Results*), deciding that we could not accurately determine between-human transmissibility based on a single observation. Thus, like our CFR analysis, our transmissibility analysis included 82 unique virus species with a total of 86 unique zoonotic transmission chains (Table S5b in *SI Data and Results*).

Lastly, we asked, *which reservoir host and virus types are associated with high death burdens in human populations?* The death count data demonstrated strong overdispersion (Figure S11 in *SI Figures*). Thus, we constructed a negative binomial GAM with the scaled observation period (i.e., number of years over which the death count was recorded) as an offset. We considered simpler Poisson GAMs, as well as zero-inflated models, but enhanced residual quantile-quantile (QQ) plots (70) suggested that these distributions fit poorly. Unlike our CFR analysis, we did not exclude viruses for which only one human case has been recorded. However, we did exclude a single virus species—Rotavirus A—for which we were unable to distinguish between deaths caused by zoonotic strains versus deaths caused by endemic human strains. Thus, our death burden models included 86 zoonotic viruses with a total of 90 transmission chains (Table S5c and S6f in *SI Data and Results*).

## Supporting information

SI Data and Results

## Supplementary Material

**SI_Data_and_Results**. Databases with variable descriptions and references, and table outputs for all selected models

**SI_Figures**. Supplementary figures (Figure S1-12)

## Additional Information

### Data and materials availability

All data, data references, code, and materials used in the analysis are publicly available in the main text, the supplementary materials, or the following github repository: https://github.com/sguth1993/zoonotic_risk_meta_analysis

## Acknowledgments

The authors thank the Boots Lab at UC Berkeley for helpful comments on this manuscript.

## Authors’ Contributions

S.G., C.E.B., D.S., and N.M. conceived the study and design. S.G., C.E.B., K.R., and N.M. collected the data and conducted the analyses. All authors participated in writing the manuscript.

