## Supplementary material for "Bats host the most virulent—but not the most dangerous—zoonotic viruses": SI Figures

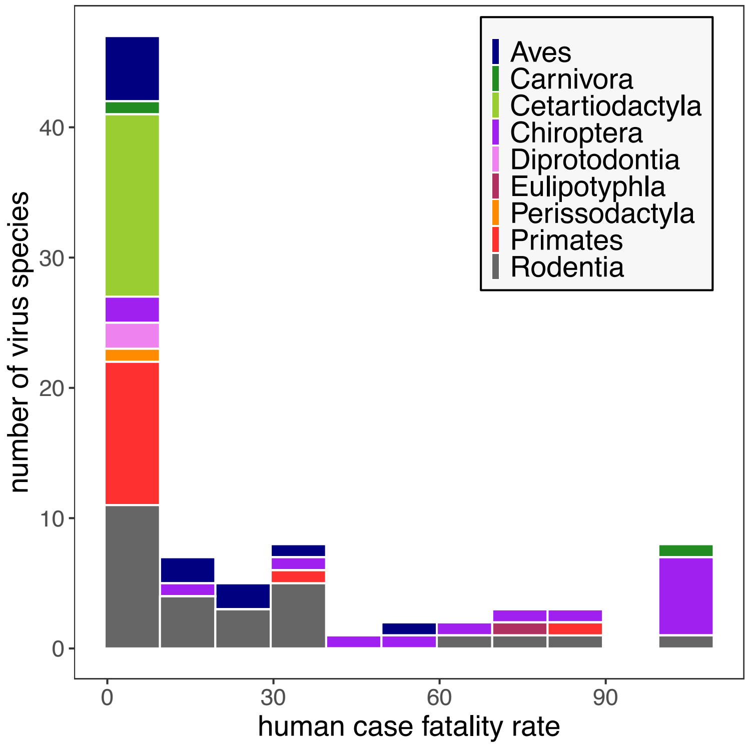


**Figure S1.** Histogram of human case fatality rates (CFRs) across all zoonotic virus species included in our virulence analysis, grouped by reservoir host type.

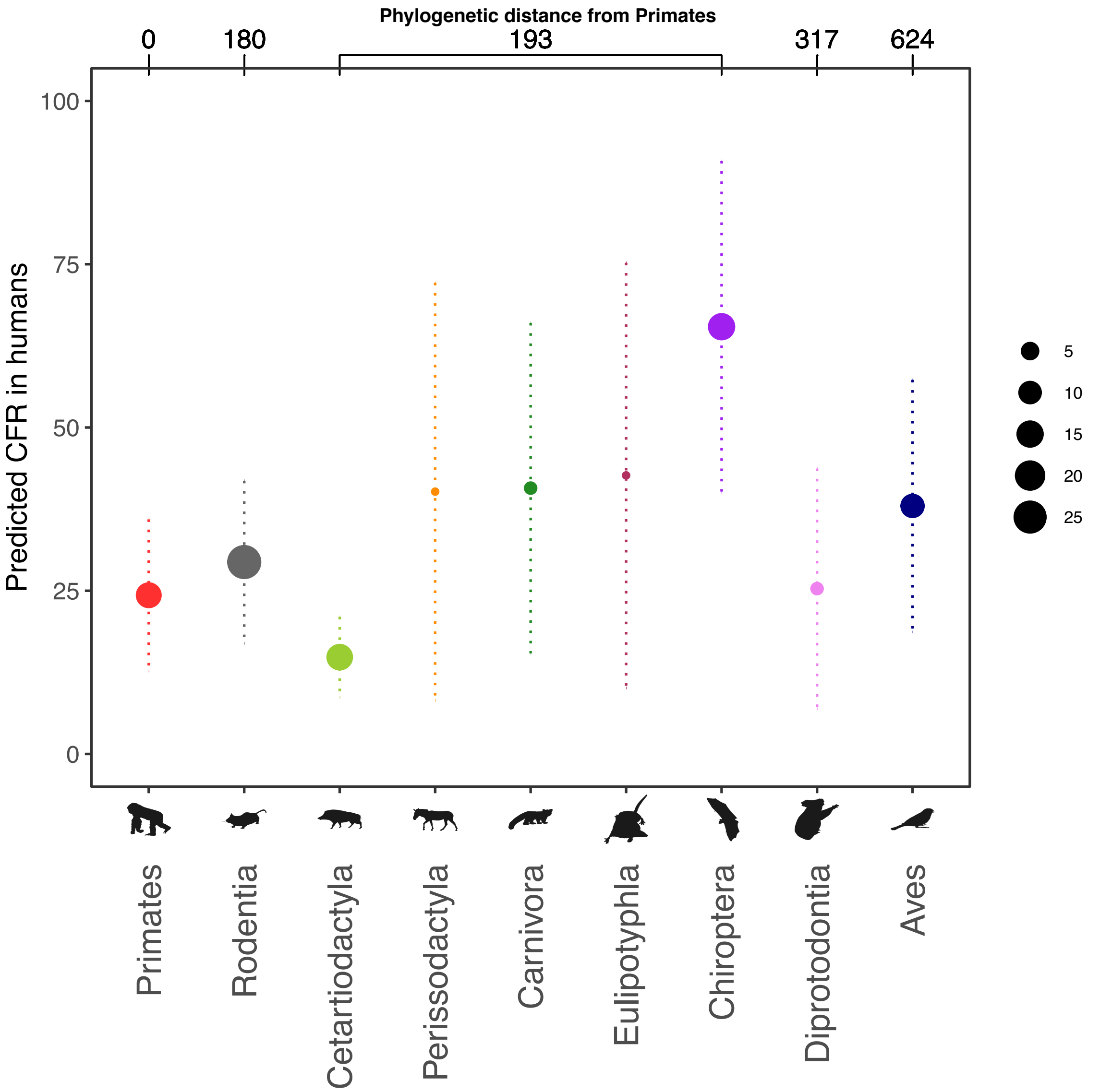


**Figure S2.** Predicted human CFR of zoonotic viruses sourced from each reservoir host group when using the top selected model of global CFR estimates.

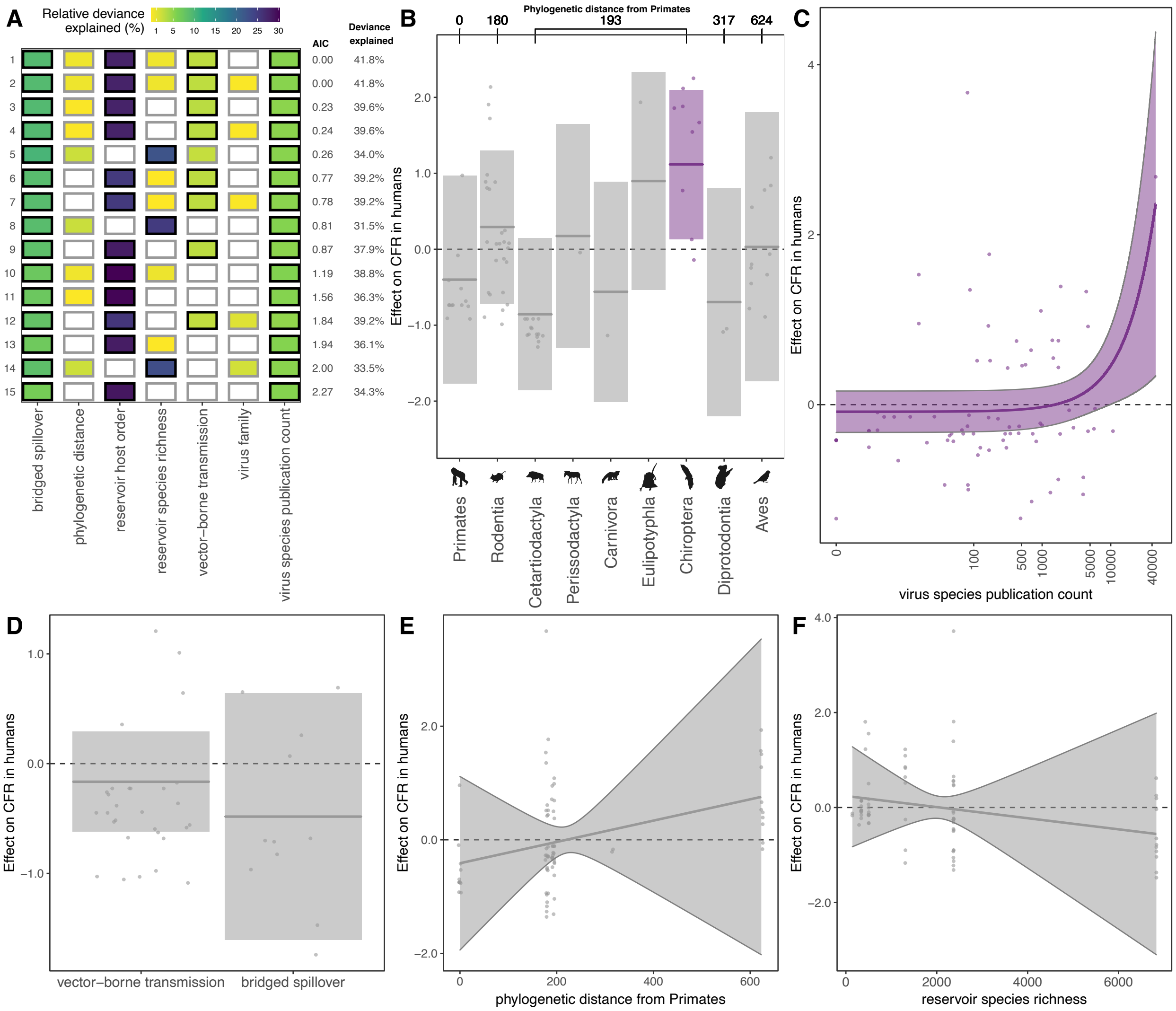


**Figure S3**. Predictors of global CFR estimates, excluding bat lyssaviruses. (A) Top 15 models ranked by AIC. Rows represent individual models and columns represent predictor variables. Cells are shaded according to the proportion of deviance explained by each predictor. Cells representing predictor variables with a p-value significance level of <0.1 are outlined in black and otherwise outlined in gray. (B-F) Effects present in the top model: reservoir host group, virus species publication count, vector-borne transmission, bridged spillover, reservoir group phylogenetic distance from Primates, and reservoir group species richness. Lines represent the predicted effect of the x-axis variable when all other variables are held at their median value (if numeric) or their mode (if categorical). Shaded regions indicate 95% CIs by standard error and points represent partial residuals. An effect is shaded in gray if the 95% CI crosses zero across the entire range of the predictor variable; in contrast, an effect is shaded in purple and considered "significant" if the 95% CI does not cross zero. Full model results are outlined in Table S6a in *SI Data and Results.* (B) Reservoir host groups are ordered by increasing cophenetic phylogenetic distance from Primates (in millions of years), as indicated on the top axis.

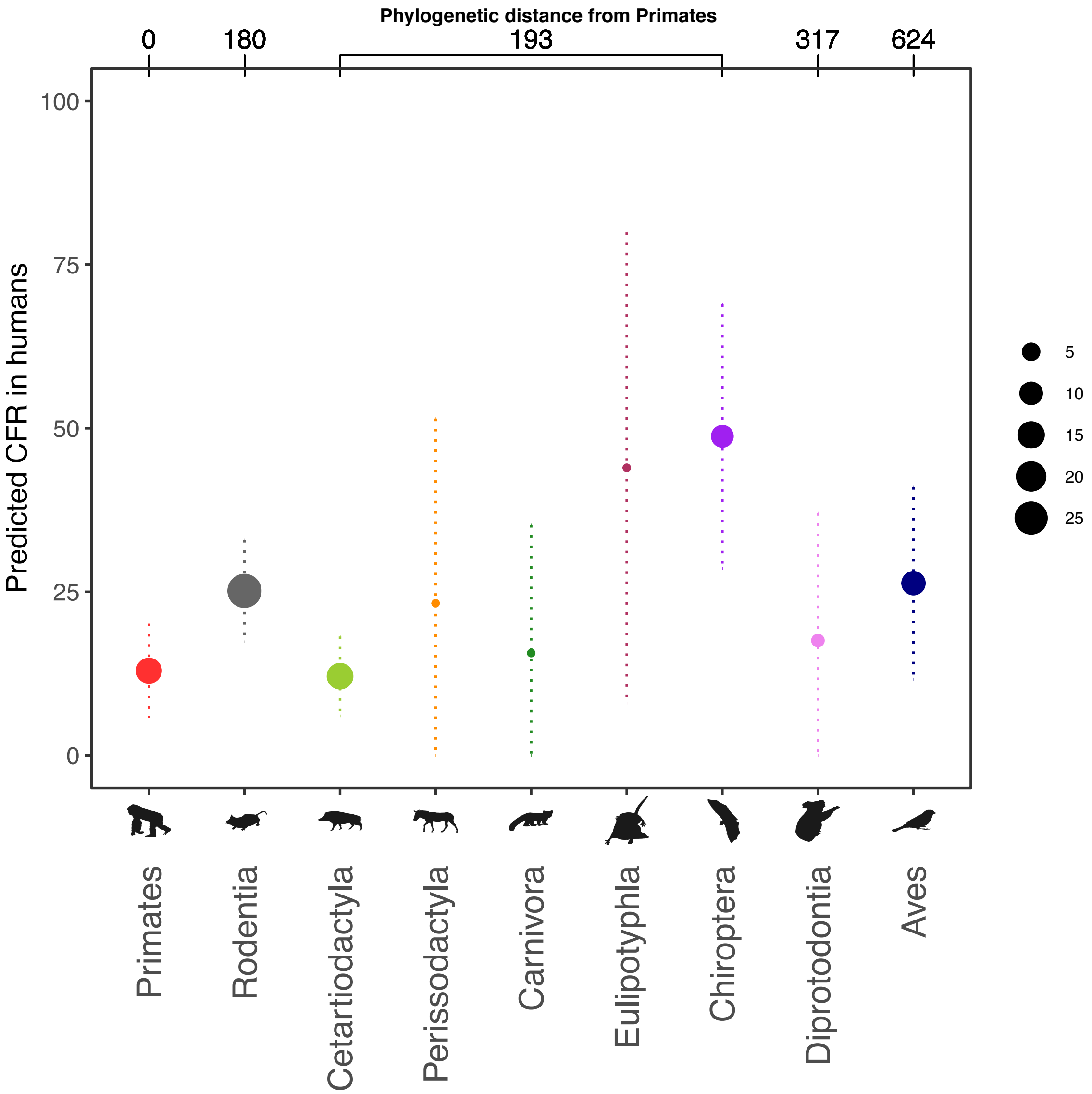


**Figure S4.** Predicted human CFR of zoonotic viruses sourced from each reservoir host group when using the top selected model of global CFR estimates, excluding bat lyssaviruses.

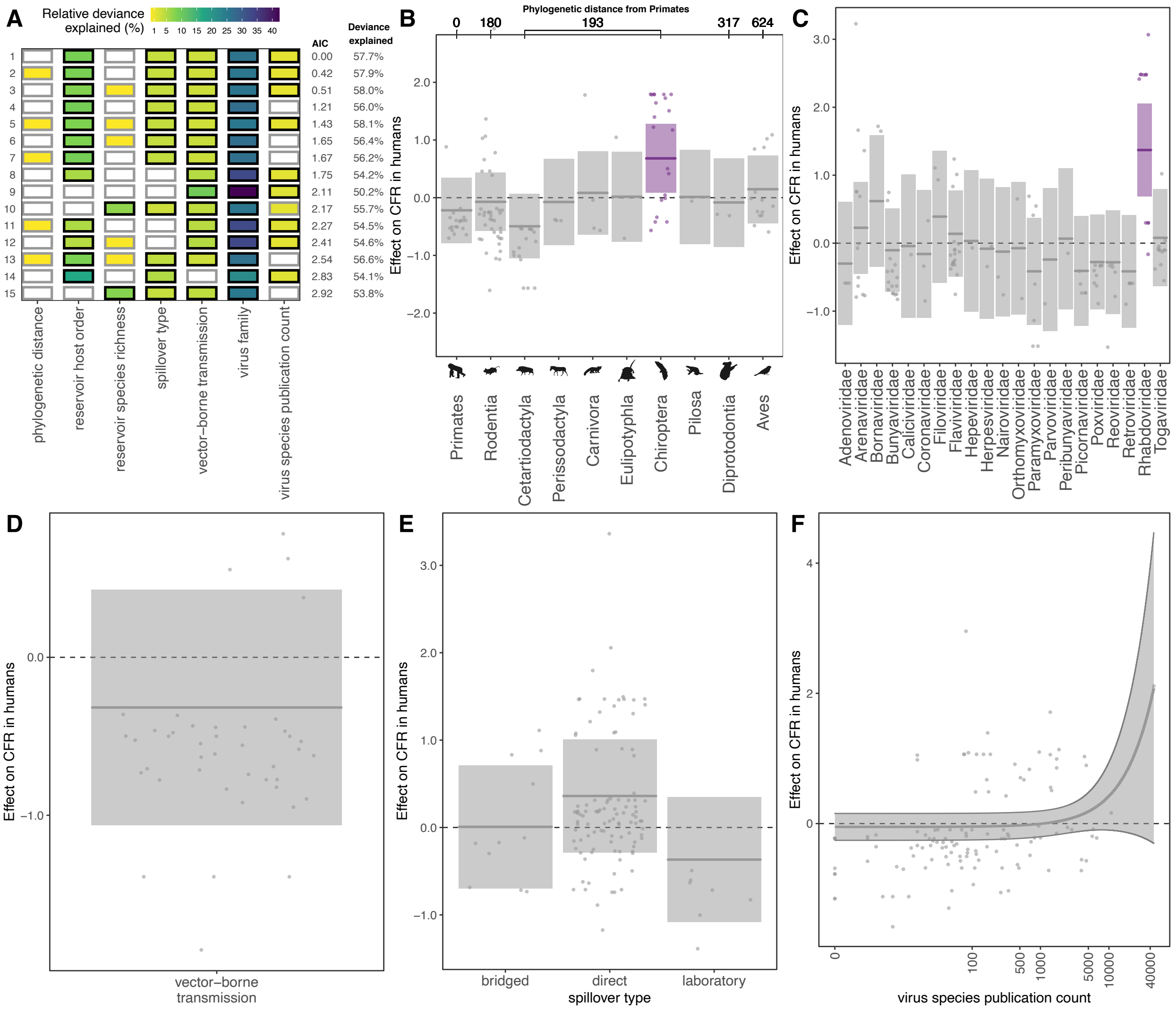


**Figure S5**. Predictors of global CFR estimates, including virus species that met a lenient definition of zoonotic. (A) Top 15 models ranked by AIC. Rows represent individual models and columns represent predictor variables. Cells are shaded according to the proportion of deviance explained by each predictor. Cells representing predictor variables with a p-value significance level of <0.1 are outlined in black and otherwise outlined in gray. (B-F) Effects present in the top model: reservoir host group, virus family, vector-borne transmission, spillover type, and virus species publication count. Lines represent the predicted effect of the x-axis variable when all other variables are held at their median value (if numeric) or their mode (if categorical). Shaded regions indicate 95% CIs by standard error and points represent partial residuals. An effect is shaded in gray if the 95% CI crosses zero across the entire range of the predictor variable; in contrast, an effect is shaded in purple and considered "significant" if the 95% CI does not cross zero. Full model results are outlined in Table S6b in *SI Data and Results.* (B) Reservoir host groups are ordered by increasing phylogenetic distance from Primates, as indicated on the top axis.

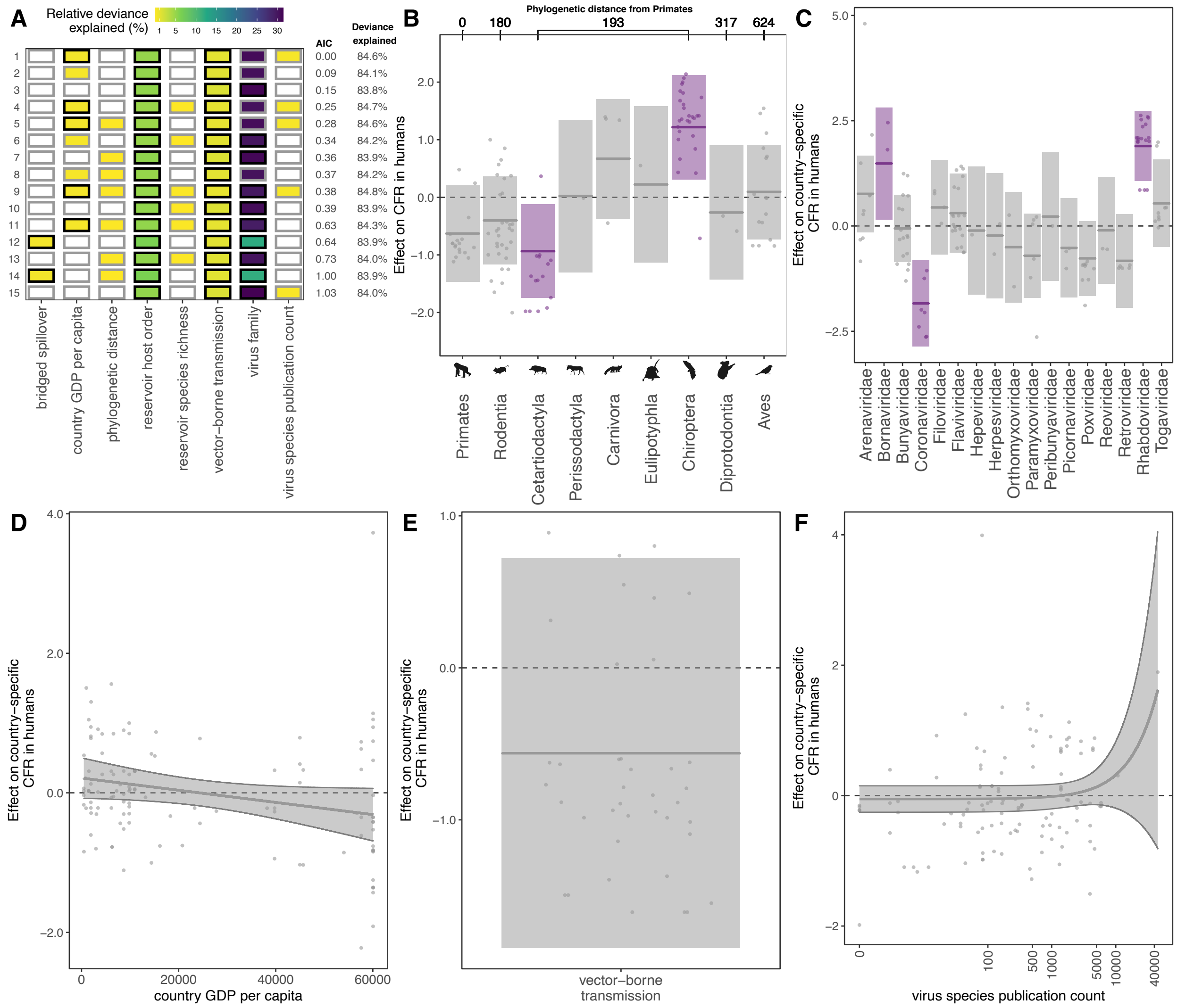


**Figure S6**. Predictors of variation among the 119 country-specific CFR estimates. (A) Top 15 models ranked by AIC. Rows represent individual models and columns represent predictor variables. Cells are shaded according to the proportion of deviance explained by each predictor. Cells representing predictor variables with a p-value significance level of <0.1 are outlined in black and otherwise outlined in gray. (B-F) Effects present in the top model: reservoir host group, virus family, country GDP per capita, vector-borne transmission, and virus species publication count. Lines represent the predicted effect of the x-axis variable when all other variables are held at their median value (if numeric) or their mode (if categorical). Shaded regions indicate 95% CIs by standard error and points represent partial residuals. An effect is shaded in gray if the 95% CI crosses zero across the entire range of the predictor variable; in contrast, an effect is shaded in purple and considered "significant" if the 95% CI does not cross zero. Full model results are outlined in Table S6c in *SI Data and Results*. (B) Reservoir host groups are ordered by increasing cophenetic phylogenetic distance from Primates (in millions of years), as indicated on the top axis.

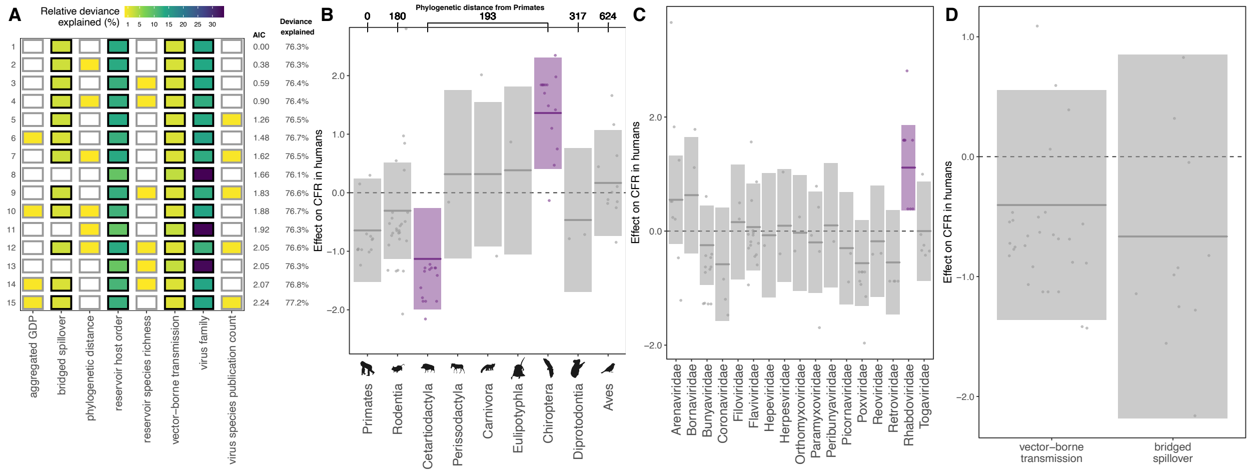


**Figure S7**. Predictors of CFRs calculated from country-level data aggregated at the level of the 86 unique zoonotic transmission chains. (A) Top 15 models ranked by AIC. Rows represent individual models and columns represent predictor variables. Cells are shaded according to the proportion of deviance explained by each predictor. Cells representing predictor variables with a p-value significance level of <0.1 are outlined in black and otherwise outlined in gray. (B-D) Effects present in the top model: reservoir host group, virus family, vector-borne transmission, and bridged spillover. Lines represent the predicted effect of the x-axis variable when all other variables are held at their median value (if numeric) or their mode (if categorical). Shaded regions indicate 95% CIs by standard error and points represent partial residuals. An effect is shaded in gray if the 95% CI crosses zero across the entire range of the predictor variable; in contrast, an effect is shaded in purple and considered "significant" if the 95% CI does not cross zero. Full model results are outlined in Table S6d in *SI Data and Results.* (B) Reservoir host groups are ordered by increasing cophenetic phylogenetic distance from Primates (in millions of years), as indicated on the top axis.

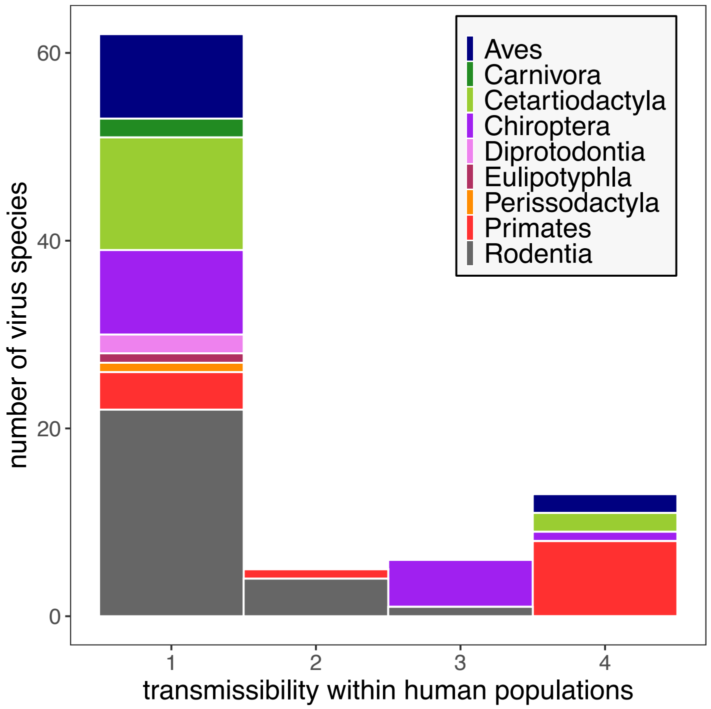


**Figure S8.** Histogram of human transmissibility rankings across all zoonotic virus species included in our analysis of capacity for forward transmission in humans, grouped by host order.

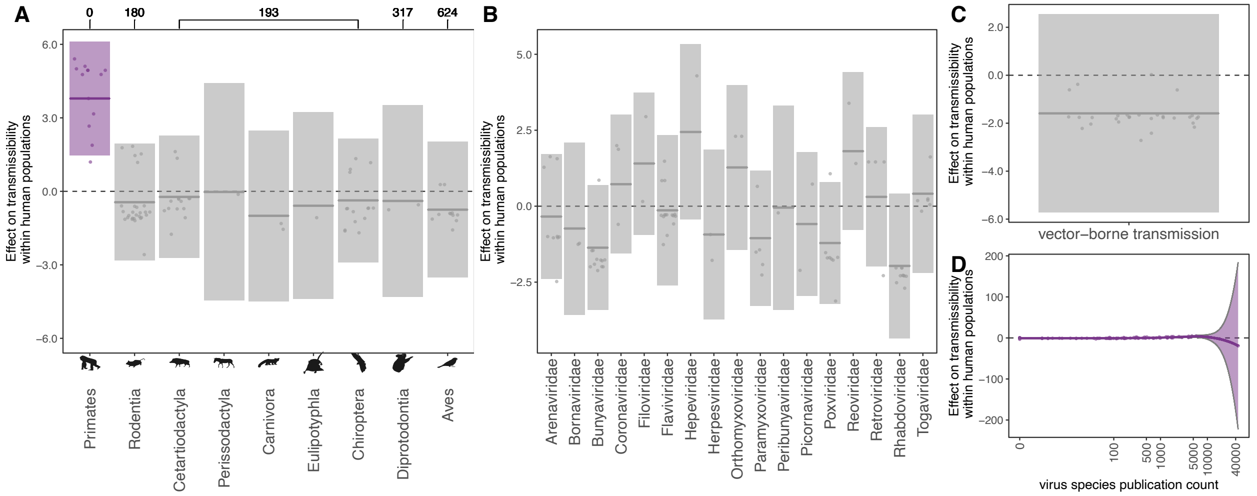


**Figure S9**. Effects present in the selected model to predict capacity for forward transmission within the human population with reservoir host order as a predictor instead of phylogenetic distance from humans. Effects include: reservoir host group, virus family, vector-borne transmission, and virus species publication count. Lines represent the predicted effect of the x-axis variable when all other variables are held at their median value (if numeric) or their mode (if categorical). Shaded regions indicate 95% CIs by standard error and points represent partial residuals. An effect is shaded in gray if the 95% CI crosses zero across the entire range of the predictor variable; in contrast, an effect is shaded in purple and considered "significant" if the 95% CI does not cross zero. Full model results are outlined in Table S6e in *SI Data and Results.*

(A) Reservoir host groups are ordered by increasing cophenetic phylogenetic distance from Primates (in millions of years), as indicated on the top axis.

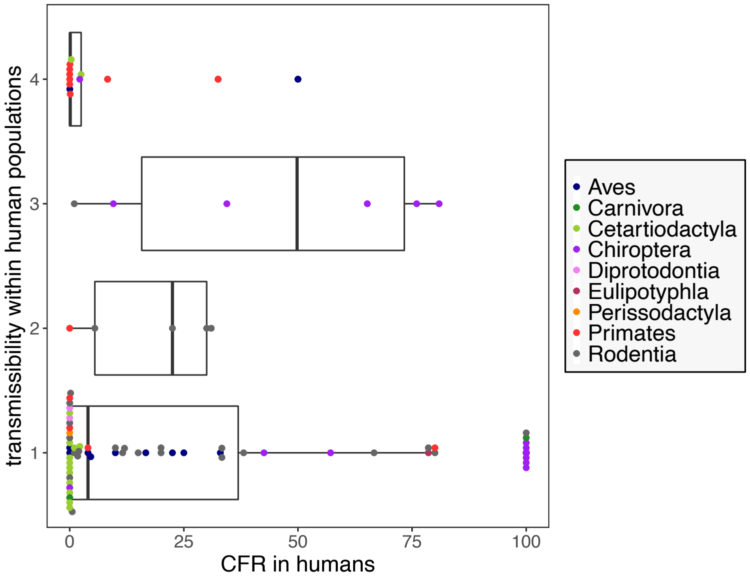


**Figure S10.** Relationship between CFR and transmissibility in humans.

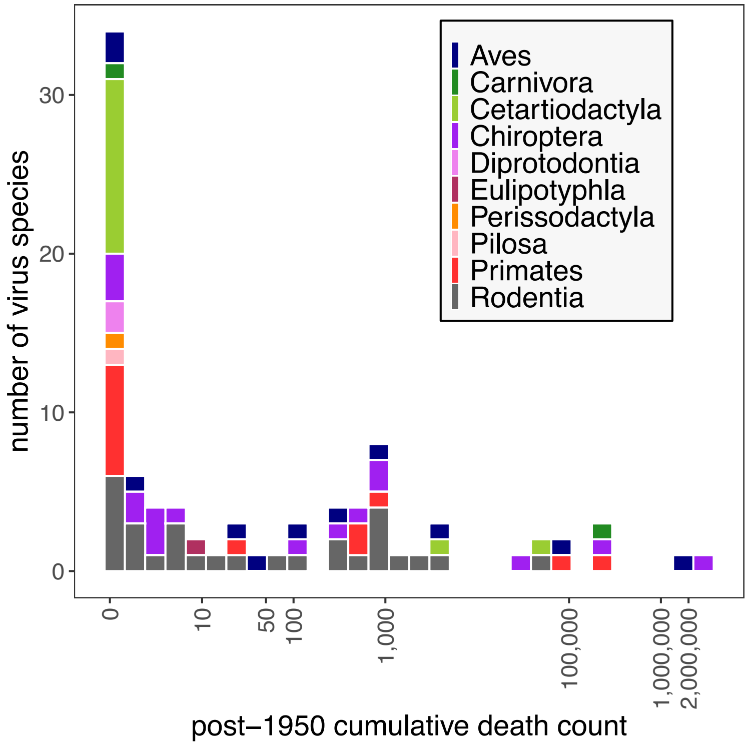


**Figure S11**. Histogram of post-1950 death counts, grouped by host order. Here, death counts were plotted to display the data distribution and are not adjusted for variation in the length of the reporting timeline. In the death burden model, we normalized counts by including an offset for the exact number of years over which deaths were recorded.

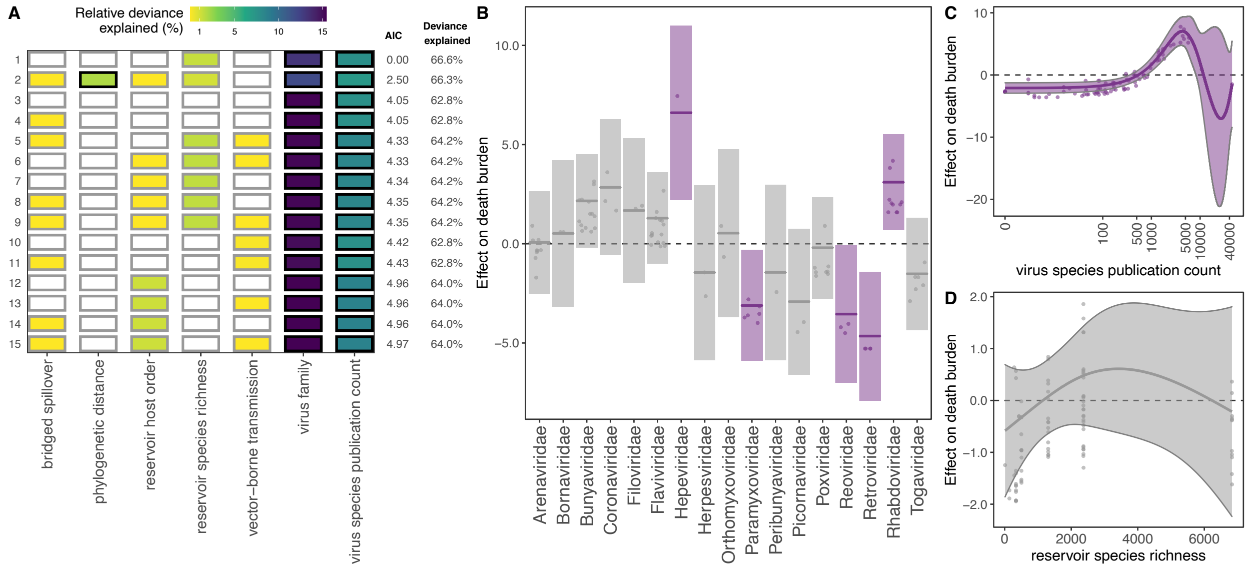


**Figure S12.** Predictors of post-1950 death burden. (A) Top 15 models ranked by AIC. Rows represent individual models and columns represent predictor variables. Cells are shaded according to the proportion of deviance explained by each predictor. Cells representing predictor variables with a p-value significance level of <0.1 are outlined in black and otherwise outlined in gray. (B-D) Effects present in the top model: virus family, virus species publication count, and reservoir group species richness. Lines represent the predicted effect of the x-axis variable when all other variables are held at their median value (if numeric) or their mode (if categorical). Shaded regions indicate 95% CIs by standard error and points represent partial residuals. An effect is shaded in gray if the 95% CI crosses zero across the entire range of the predictor variable; in contrast, an effect is shaded in purple and considered "significant" if the 95% CI does not cross zero. Full model results are outlined in Table S6f in *SI Data and Results.* (C) Reservoir host groups are ordered by increasing cophenetic phylogenetic distance from Primates (in millions of years), as indicated on the top axis.
